# Rapidly Assessing the Quality of Targeted Proteomics Experiments Through Monitoring Stable-isotope Labeled Standards

**DOI:** 10.1101/409805

**Authors:** Bryson C. Gibbons, Thomas L. Fillmore, Yuqian Gao, Ronald J. Moore, Tao Liu, Ernesto S. Nakayasu, Thomas O. Metz, Samuel H. Payne

## Abstract

Targeted proteomics experiments based on selected reaction monitoring (SRM) have gained wide adoption in clinical biomarker, cellular modeling and numerous other biological experiments due to their highly accurate and reproducible quantification. The quantitative accuracy in targeted proteomics experiments is reliant on the stable-isotope, heavy-labeled peptide standards which are spiked into a sample and used as a reference when calculating the abundance of endogenous peptides. Therefore, the quality of measurement for these standards is a critical factor in determining whether data acquisition was successful. With improved MS instrumentation that enables the monitoring of hundreds of peptides in hundreds to thousands of samples, quality assessment is increasingly important and cannot be performed manually. We present Q4SRM, a software tool that rapidly checks the signal from all heavy labeled peptides and flags those that fail quality control metrics. Using four metrics, the tool detects problems both with individual SRM transitions and the collective group of transitions that monitor a single peptide. The program’s speed enables its use at the point of data acquisition and can be ideally run immediately upon the completion of an LC-SRM-MS analysis.

## Introduction

Selected reaction monitoring (SRM), also known as multiple reaction monitoring (MRM), is a data acquisition technique used in targeted analysis of molecules, including targeted proteomic studies. It exploits the unique capability of triple quadrupole (QQQ) mass spectrometers to monitor the predefined precursor/fragment ion pairs of specific molecules of interest throughout a liquid chromatography (LC) elution profile. Compared to shotgun proteomics, targeted proteomics using SRM has high selectivity, high sensitivity and wide linear dynamic range^1^–^3^, which makes it especially useful in the accurate and reproducible quantification of low abundance proteins in highly complex biological samples. SRM has been widely used in the fields of biomarker discovery^4^–^7^, analysis of protein post-translational modifications^8^ and characterization of biological protein networks^4,9^.

In the recent years, multiple technical advantages have greatly improved the throughput of SRM analyses, allowing for the quantification of hundreds of peptides in a single analysis^6,7,10^. For example, a single 800-plex SRM assay (e.g. 400 unlabeled and heavy labeled peptide pairs and 2400 transitions with retention time scheduling) using ultra-high performance liquid chromatography (UHPLC) has been developed to quantify proteins in plasma^11^. Additionally, advanced labeling techniques utilizing in vitro proteome–assisted MRM for protein absolute quantification (iMPAQT) demonstrated the capability of SRM in genome-wide protein quantification of over 18,000 human proteins^12^. Moreover, the scan speed of triple quadrupole instrumentation has been greatly improved in recent releases of commercial instrumentation. The newly developed TSQ Altis (released in 2017) can scan more than 600 transitions per second, which is 6 times more than a traditional QQQ scan speed of 100 transitions per second. The breadth of measurement enabled by these technological improvements to QQQ mass spectrometry has increased the feasibility and popularity of large scale targeted proteomics studies, a major application of which will be in clinical studies where up to hundreds of protein candidates need to be quantified in hundreds of clinical specimens^13^.

Quantitative accuracy is a primary motivating factor for utilizing a targeted proteomics protocol. The precise and reproducible absolute quantification produced by SRM assays is essential to many clinical and laboratory experiments^14,15^. As the abundance of an endogenous peptide is calculated from the measurement of the spiked-in reference standard, it is essential to assess the data quality of these references^16^. In early applications of targeted proteomics, when instrument speed greatly limited the number of transitions that could be monitored, much of this quality assessment was done manually. However, recent improvements in instrument performance and experimental design have enabled a dramatic increase in the number of target peptides and associated SRM transitions, which makes manual quality assessment an untenable and laborious task.

A variety of computational tools assist in SRM experiment design and data analysis. The first task in creating an SRM experiment is the choice of proteins and representative peptides to monitor. Achieving a reliable protein abundance requires appropriately choosing peptides that have a strong signal and are free from interferences in the biological matrix. Numerous computational tools exist to facilitate assay design by identifying peptides and refining SRM transitions^17^–^20^. To help share these assays and eliminate time spent designing the same transitions at multiple institutions, community portals have begun to host well-designed and vetted assays^21^–^23^. Analyzing the experimental data requires significant computational effort to align files across replicates and experimental conditions, pick peaks and produce quantitative values, normalize data and perform statistical tests, etc.^24^–^26^.

Among the many tools that are used in the SRM community, there remains a need for a tool to assist in quality assessment. In particular, the rapid quality assessment of reference transitions immediately following data acquisition lacks an easy-to-use tool. Therefore, we have created Q4SRM, a software tool that rapidly checks the signal from all heavy-labeled target peptides and flags those that fail quality control metrics.

## Methods and implementation

Q4SRM is a C# .NET application designed to perform quality assessment of transitions associated with the heavy-labeled reference peptides that are spiked into a sample. Because these peptides are spiked into every sample, their transitions are expected to be easily identified in each MS result file. The software is open source under the BSD license and available on GitHub at: https://github.com/PNNL-Comp-Mass-Spec/Q4SRM. The software expects two types of input. The first is a Thermo .RAW file representing the data acquisition from a triple quadrupole instrument, e.g. TSQ Vantage or TSQ Altis. To read this file format, the software utilizes the I/O codex that is part of the RawFileReader NuGet package distributed by Thermo (San Jose, CA); these DLLs are included with the Q4SRM executable. The second input is a user generated file which contains cutoffs and thresholds used in determining which data points are flagged with warnings. This is a simple tab-delimited text file where each row describes thresholds to be used for a specific peptide. An example file is available in the project’s GitHub repository. If one desires uniform thresholds for all peptides, this file can be omitted and the thresholds specified directly in the interface.

To identify the transitions for reference/heavy peptides, the program looks for a keyword in the “Name” (TSQ Vantage) or “Compound Name” (TSQ Altis) field of the SRM Table (contained in the Instrument Method portion of the .RAW file); transitions lacking the keyword are ignored. It is customary in our lab to name the transitions associated with heavy peptides with the string “heavy” or “hvy”, e.g. ‘VSGVATDIQALK_heavy’. Note that multiple transitions for the same peptide have the same name. Since the program is open source, it is possible to adapt this parsing step for other keywords, if different conventions are used in other laboratories. The SRM Table portion of the .RAW file also provides a parent/precursor *m/z*, product *m/z*, and a start and stop time (in minutes) for each transition. The heavy transitions are then grouped according to name, so that all transitions for each precursor can be associated appropriately in the output.

### Data extraction and metrics.

For each heavy transition, we gather four pieces of information from the .RAW file: max intensity, time of max intensity, median intensity, and the sum total intensity during the scheduling window. With these pieces of information, we compute four metrics.

Two metrics are computed based on information relating to a single transition. The first metric, called *peak position*, calculates the time between the start or stop of transition acquisition and the time of the max intensity. The user defined cutoff (floating point number representing time in minutes) dictates what is considered an acceptable minimal value. The reason for this metric is to ensure that peaks fully elute within their expected scheduled time, and are not clipped or truncated, which in part may be due to degradation of the LC column performance. Internally, the software scales the input values (time in minutes) to unit distance considering the entire time of a run in the range 0-1, analogous to the Normalized Elution Time strategy^27^, however this is not exposed to the user. The second metric, called *S/N heuristic*, calculates the ratio of the maximal intensity to the median intensity. For the purpose of calculating the median value, all values below 5 are excluded. A user defined threshold (floating point number) dictates what is considered an acceptable minimal value. This approximates the signal-to-noise as the intensity of the transition relative to the background intensity of unrelated signal. We recognize that there are many ways to calculate a S/N ratio, and this heuristic is not intended to be a thorough calculation (which would involve a more statistical characterization of the noise). This heuristic is designed to quickly assess whether there is a strong and distinct peak relative to the other intensities within the acquisition window.

The last two metrics are computed based on information relating to all transitions belonging to the same peptide, i.e. a group of transitions. The first metric, called *Total signal*, is the sum of the intensities for all transitions in the group. The user defined threshold (integer number) dictates what is considered an acceptable minimum value. The reason for this metric is to ensure that sufficient signal exists for all transitions in the group, which is required in order to have an accurate quantitative measurement. The second metric, called *Peak concurrence*, calculates the difference between the time of the max intensity for each transition in the group, providing a warning for situations where the transitions do not have a reasonably concurrent max intensity. Again, users specify a threshold (time in minutes) that defaults to 0.5 minutes or 30 seconds. It is expected that the time of max intensity of transitions for the same peptide should be identical; however, an interference in one transition may cause these values to be out of sync.

The program produces both text and graphical output. The text output file is a tab-delimited text file that contains the four computed metrics as well as other associated information for each peptide. An example of the output is in the GitHub repository. For graphical output, the program produces an image of each group of transitions, similar to what is seen in Figures 1 and 2. There is also a summary image that shows which data points give warnings and which pass QC.

**Figure 1.**
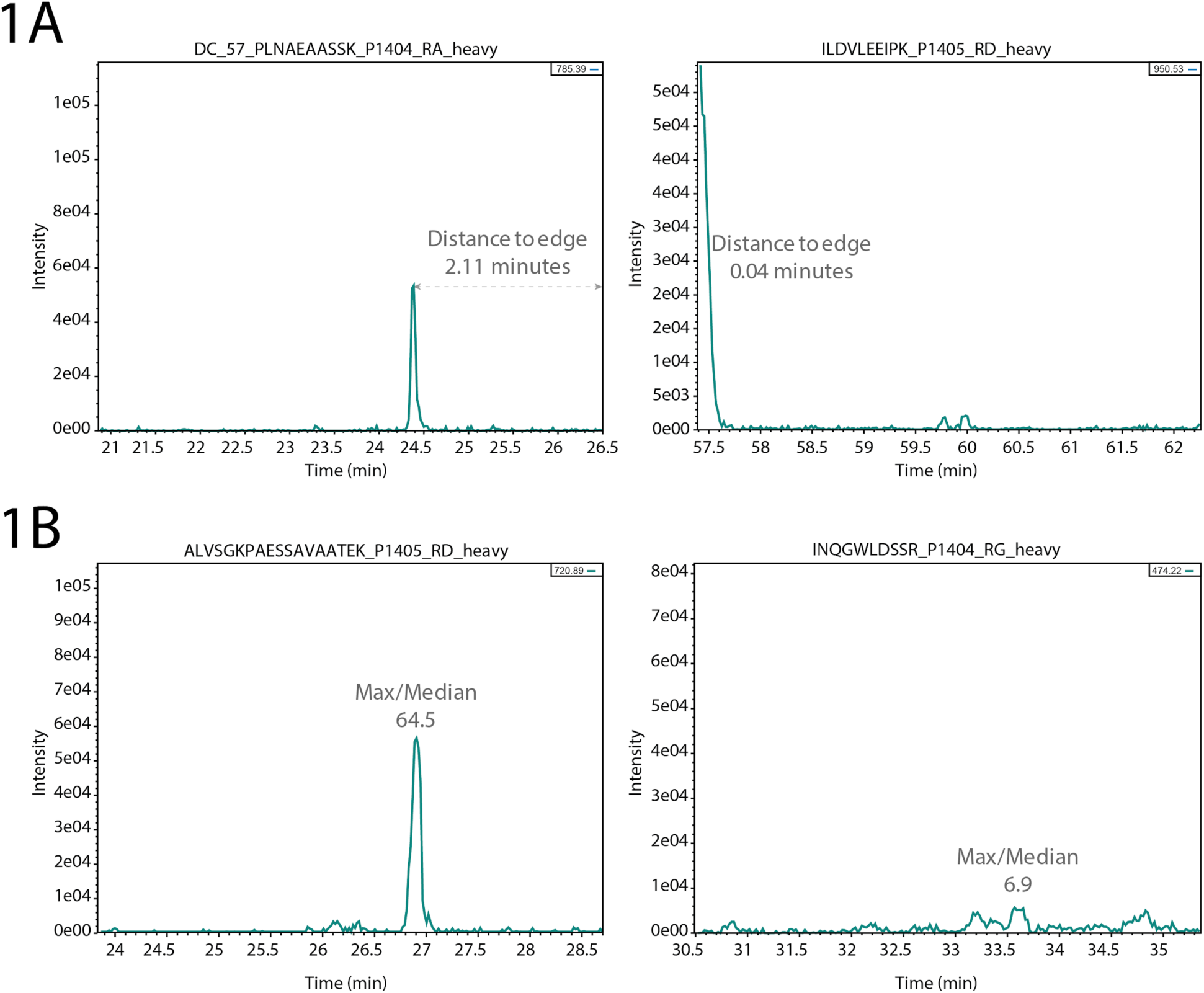
Single transition metrics. A) The distance of a peak to the edge of the scheduled time window indicates whether the peak was potentially incompletely measured (right). B) The ratio of the maximal intensity to the median indicates whether the transition has a strong intensity relative to the background signal. Low ratios (right) suggest a poorly defined signal and may not be reliable when used as a reference to measure endogenous peptide abundance.

**Figure 2.**
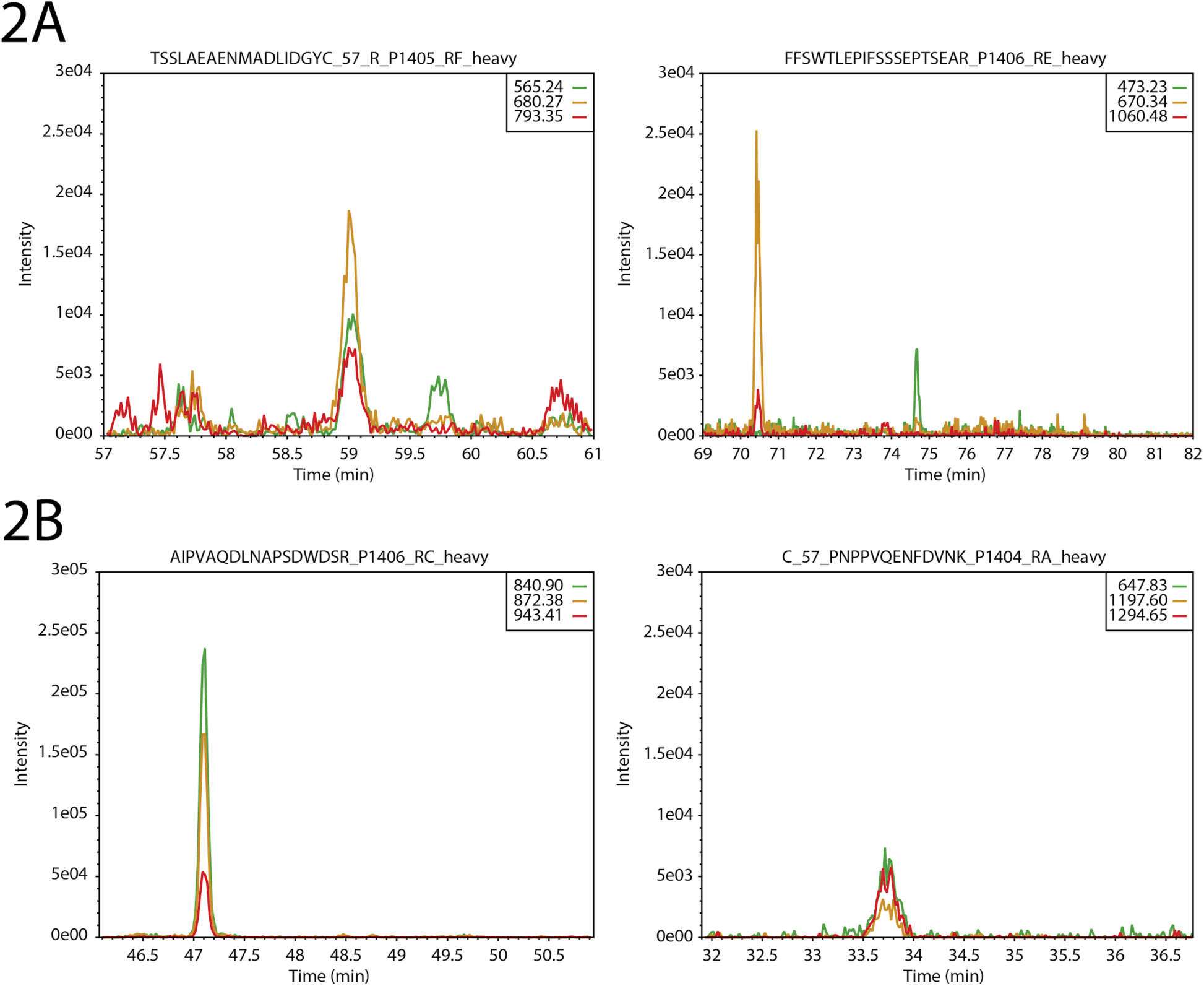
Multi-transition metrics. A) Transitions that belong to the same peptide should have identical elution profiles. On the left is shown a set of transitions that all elute at the same time; on the right we see that the peak of the yellow and red transition peaks (∼70 min) are distant from the green transition peak (∼74 min). B). Transitions should have sufficient intensity as a group. On the left is a strong and intense set of transitions (note y-axis scale); on the right is a set of low intensity transitions. Depending on the user-defined threshold for the total intensity metric, this set of transitions may yield a warning.

Q4SRM is available with both a graphical user interface and a command-line interface. The graphical user interface version facilitates the selection of input files and the adjustment of settings, and also provides a viewer mode where the user can view the summary plot and get details on the different points. The command-line version provides access to the same settings as the graphical user interface version and facilitates use of Q4SRM with computational pipelines. A pictorial user guide that walks through download and use of both user interfaces is included on the GitHub repository wiki page (https://github.com/PNNL-Comp-Mass-Spec/Q4SRM/wiki).

## Results

Targeted proteomics experiments are rapidly growing in their capacity to measure a large number of peptide targets. Although a full analysis of the data will happen in the days and weeks that follow data acquisition and in the context of the entire experiment, it is essential to rapidly assess the quality of the data immediately as it is generated to determine whether the run was successful. To assist in point-of-acquisition quality assessment of LC-SRM-MS datasets, we have created the Q4SRM software package. This easy-to-use package rapidly checks transitions for each heavy-labeled peptide against a suite of essential QC metrics, and provides simple and interpretable output to the operator, including a list of flagged transitions that need further manual inspection. Q4SRM is a lightweight software tool that can be installed on the computers that control MS instrumentation. Even for files with thousands of scheduled transitions, the software takes less than 1 minute to analyze a single Thermo .RAW file.

Two metrics assessed by Q4SRM report information on individual transitions (Figure 1). First, the program measures the distance from the maximal peak intensity to the edge of the scheduled acquisition window. This metric flags a transition with a warning when the peak maximum is too close to the edge of the window, signaling that this peak is potentially clipped (Figure 1A). This would mean that the quantitation will not be accurate because some of the peptide’s elution profile was not measured. The user specified threshold should be set in relation to the schedule window size, the expected LC peak widths and the operator’s personal tolerance. The second metric derived from data for a single transition is an approximate measure of signal to noise (Figure 1B). Although S/N may be calculated a variety of ways, our goal here is to quickly determine whether there is a problem with the data. Therefore, the S/N heuristic calculated by Q4SRM is the ratio between the peak maximum intensity and the median intensity. For example, a value of 10 means that the peak maximum is 10 times greater than the median intensity during the schedule window, thus indicating that the peak is strongly intense above background signal. With these two metrics, users can be confident that the measured transition of the heavy labeled standard is both clearly within the scheduled LC time window and sufficiently intense to serve as a reference in the calculation of an accurate quantitative value for the endogenous peptide.

Two metrics assessed by Q4SRM report information about the group of transitions related to the same peptide (Figure 2). Despite good performance of individual transitions, it is necessary that the group performs as expected. The first metric measures how close in elution the transitions are to each other, or peak concurrence. Since each transition is intended to measure the same peptide, they are expected to have an identical elution profile. However, due to potential interferences or missing signal, the transitions may appear out of sync with each other. Figure 2A shows two sets of transitions. One transition has acceptable peak concurrence despite being low abundance (Figure 2A, left); in the other set of transitions, one of the peaks is clearly several minutes after the elution of the other two (Figure 2A, right). The second metric, total ion intensity, simply measures the total intensity of all transitions associated with a peptide (Figure 2B). This metric can be set to a different threshold for each peptide, since each peptide and transition is expected to have a different characteristic response during the LC-SRM-MS analysis. A convenient method for setting these values is to take the intensity values from a data acquisition when the peptides were spiked into either a blank background or a sample matrix. By averaging values over several initial testing runs, a threshold can be set that is appropriate for the observed range of response. Failures of this metric can signal a few different challenges. First, there might be a problem with the spike-in level during sample preparation. Second, there might be an instrument performance problem causing low signal. Finally, it is possible that the peptide was completely out of range of the schedule window (due to LC column problems).

## Conclusions

Before data analysis begins in earnest, assessing the quality of the acquired data is essential. For experiments that contain many samples, and where the time between data acquisition and data analysis is long, this QC/QA step should not be delayed; rather QC/QA should happen immediately upon data acquisition to give feedback as soon as possible to the instrument operator^28^. To fill this need in targeted proteomics studies, we created Q4SRM which can analyze the heavy labeled reference peptides in an LC-SRM-MS data file within one minute. It quickly computes a set of four essential QC metrics that helps to identify low quality SRM transitions. The number of flagged transitions for any dataset depends on user-specified thresholds and instrument performance. We have found it to be an essential tool for maintaining high data quality and instrument health. To assist in long-term monitoring of QC metrics, Q4SRM’s text output contains the metrics on all transitions. With this information, users can collate and compare output across many different acquisition files using data analytic platforms like R or Jupyter.

## Author Contributions

BCG, TLF, ESN and TOM conceived project; BCG implemented software; TLF acquired mass spectrometry data; TLF, BCG, YG, and SHP analyzed data; RJM, TL, ESN and TOM provided scientific leadership and oversight; BCG, YG and SHP wrote the manuscript with input from all authors.

### Acknowledgements

We thank Geremy Clair for assistance with figures. This work was supported by the NIH National Institute of General Medical Sciences grant P41GM103493 (Richard Smith, PNNL), the National Cancer Institute Clinical Proteomic Tumor Analysis Consortium (CPTAC) grant 1U24CA210972 (S.H.P.) and 5U24CA210955 (T.L.), the NCI Early Detection Research Network (EDRN) Interagency Agreement ACN15006-001 (T.L.), the MoTrPAC consortium U24DK112349 (Joshua Adkins, PNNL). Additional support was provided by the TEDDY Study Group, which is funded by the National Institute of Diabetes and Digestive and Kidney Diseases (NIDDK), National Institute of Allergy and Infectious Diseases (NIAID), National Institute of Child Health and Human Development (NICHD), National Institute of Environmental Health Sciences (NIEHS), Centers for Disease Control and Prevention (CDC), Juvenile Diabetes Research Foundation (JDRF), and supported in part by the National Center for Advancing Translational Sciences Clinical and Translational Science (NCATS) Awards to the University of Florida and the University of Colorado. Work was performed in the Environmental Molecular Science Laboratory, a U.S. Department of Energy (DOE) national scientific user facility at Pacific Northwest National Laboratory (PNNL) in Richland, WA. Battelle operates PNNL for the DOE under contract DE-AC05-76RLO01830.

